# Neurexin 3 is Essential for the Specific Wiring of a Color Pathway in the Mammalian Retina

**DOI:** 10.1101/2023.02.13.527055

**Authors:** Vincent P Kunze, Juan M Angueyra, John M Ball, Michael B Thomsen, Xiaoyi Li, Adit Sabnis, Francisco M Nadal-Nicolás, Wei Li

## Abstract

Precise wiring within sensory systems is critical for the accurate transmission of information. In the visual system, S-cone photoreceptors specialize in detecting short-wavelength light, crucial to color perception and environmental cue detection. S-cones form specific synapses with S-cone bipolar cells (SCBCs), a connection that is remarkably consistent across species. Yet, the molecular mechanisms guiding this specificity remain unexplored. To address this, we used the cone-dominant ground squirrel for deep-sequencing of cone subtype transcriptomes and identified *Nrxn3* as an essential molecule for the S-cone to SCBC synapse. Using transgenic mouse models, we further examined the role of *Nrxn3* in S-cones and discovered a significant reduction of SCBC connections in the absence of *Nrxn3*. This finding extends the known functions of neurexins, typically associated with synapse regulation, by highlighting their essential role in a specific synaptic connection for the first time. Moreover, the differentially expressed genes identified here pave the way for further investigations into the unique functions of cone subtypes.

## Introduction

Understanding the mechanisms for normal neuronal connection establishment and wiring is crucial for addressing neural degenerative diseases and for developing effective treatments aimed at restoring neural wiring and function. Precise wiring is essential in sensory systems to ensure accurate information relay to higher brain regions. Due to its structured, well-documented architecture and accessibility, the neural retina serves as an exemplary model for investigating synapse formation and circuit organization principles (*1*).

In the mammalian retina, vision is initiated with two types of photoreceptors, rods and cones, which are responsible for light detection. Rods are operational in low-light conditions, such as moonlight, whereas cones function in daylight and are involved in color detection through various cone subtypes. Typically, most mammals are dichromats, possessing two cone subtypes: M-cones (medium wavelength-sensitive) and S-cones (short wavelength-sensitive). These cone subtypes contain distinct opsin proteins that are sensitive to specific light wavelengths, enabling downstream neurons to create color-contrast codes. S-cones utilize a short wavelength-sensitive opsin 1 (SWS1) pigment, with peak absorption wavelengths (λ_max_) ranging from ultraviolet sensitivity (350–370 nm, as seen in murid rodents and bats) to blue sensitivity (400–457 nm, observed in primates, cats, and dogs) (*2, 3*). The retinal pathway carrying S-cone signals appears to be ancient in evolutionary sense (*4*). In particular, S-cones form specialized connections with S-cone bipolar cells (SCBCs), a cell type that is highly conserved among mammals (*5–10*). In turn, SCBCs exclusively receive input from S-cones, which typically constitute a minor fraction (<10%) of the total cone population (*11–13*). Therefore, the accurate wiring of S-cones and SCBCs is crucial for the transmission of short wavelength signals. However, the mechanisms responsible for the establishment and upkeep of these connections remain elusive, underscoring a significant gap in our understanding of retinal circuitry.

To uncover the mechanisms behind the specific synaptic connections of S-cones, we analyzed the transcriptome of retinal photoreceptors in the thirteen-lined ground squirrel (TLGS). Our aim was to identify differentially expressed genes (DEGs) associated with cell adhesion molecules that could differentiate between cone subtypes. The choice of TLGS retinas was strategic due to their high cone density (∼85%) and the absence of mixed cones, which are prevalent in mice (85% cones in the mouse retina express both S- and M-opsin) and complicate analyses (*11, 13–15*). Although previous research has employed droplet-based high-throughput sequencing to profile the retinas of various species (*16–19*), these methods have not yet successfully pinpointed DEGs specific to S-cone wiring. This limitation could stem from the relatively shallow sequencing depth of droplet-based techniques, potentially missing genes with low expression levels (*20*). To address this, we adopted a targeted deep-sequencing strategy (SMART-seq) for individual photoreceptors, leading to the discovery of S-cone-specific expression of *Nrxn3*. To further validate these findings and investigate *Nrxn3*’s function, we examined cone-specific *Nrxn3*-knockout mice. These mice exhibited a notable reduction in S-cone to SCBC synapses compared to controls, underscoring *Nrxn3*’s crucial role in synapse formation and/or maintenance.

Our study highlights the strength of manual cell collection combined with the SMART-seq method in unveiling new molecular mechanisms that govern neuronal connections at a specific synapse of interest. It represents the first example of *Nrxn3*’s role in a specific neural circuitry. The discovery also provides a molecular target vital for reestablishing proper cone connections in future treatments of retinal degenerative diseases.

## Results

### Deep sequencing of manually collected TLGS photoreceptors generated high-quality genetic profiles of cone subtypes

To identify molecules responsible for cone-subtype specific synaptic wiring, we turned to sequencing of cones in the TLGS retina, where cones are highly abundant and S- and M-cone types are clearly distinguished by the opsin they express; i.e., cones do not co-express different opsins in the same cell. High-throughput single cell RNA-seq applications are a common tool to rapidly investigate thousands of cells for gene expression, but sequencing depth per cell remains low. Instead, we applied the SMART-seq approach that sequences a smaller number of isolated cells, with the advantage of receiving more reads per cell of interest, resulting in a higher gene detection rate (*20*). Hence, we developed a protocol to dissociate and isolate live S- and M-cone photoreceptors (*21*). We took advantage of the fact that cone outer segment disks are open to the extracellular space by utilizing an anti-S-opsin antibody directed at the extracellular N-terminal domain (Fig. 1 A). Antibody labeling in a live whole-mount tissue preparation resembles the mosaic visible in the fixed tissue preparation (Fig. 1 B-C). This approach enabled us to identify live S-cones following tissue separation, using a micromanipulator to collect and sort fluorescent S-cones and non-fluorescent M-cones for targeted sequencing (Figure 1 D-F).

**Figure 1.**
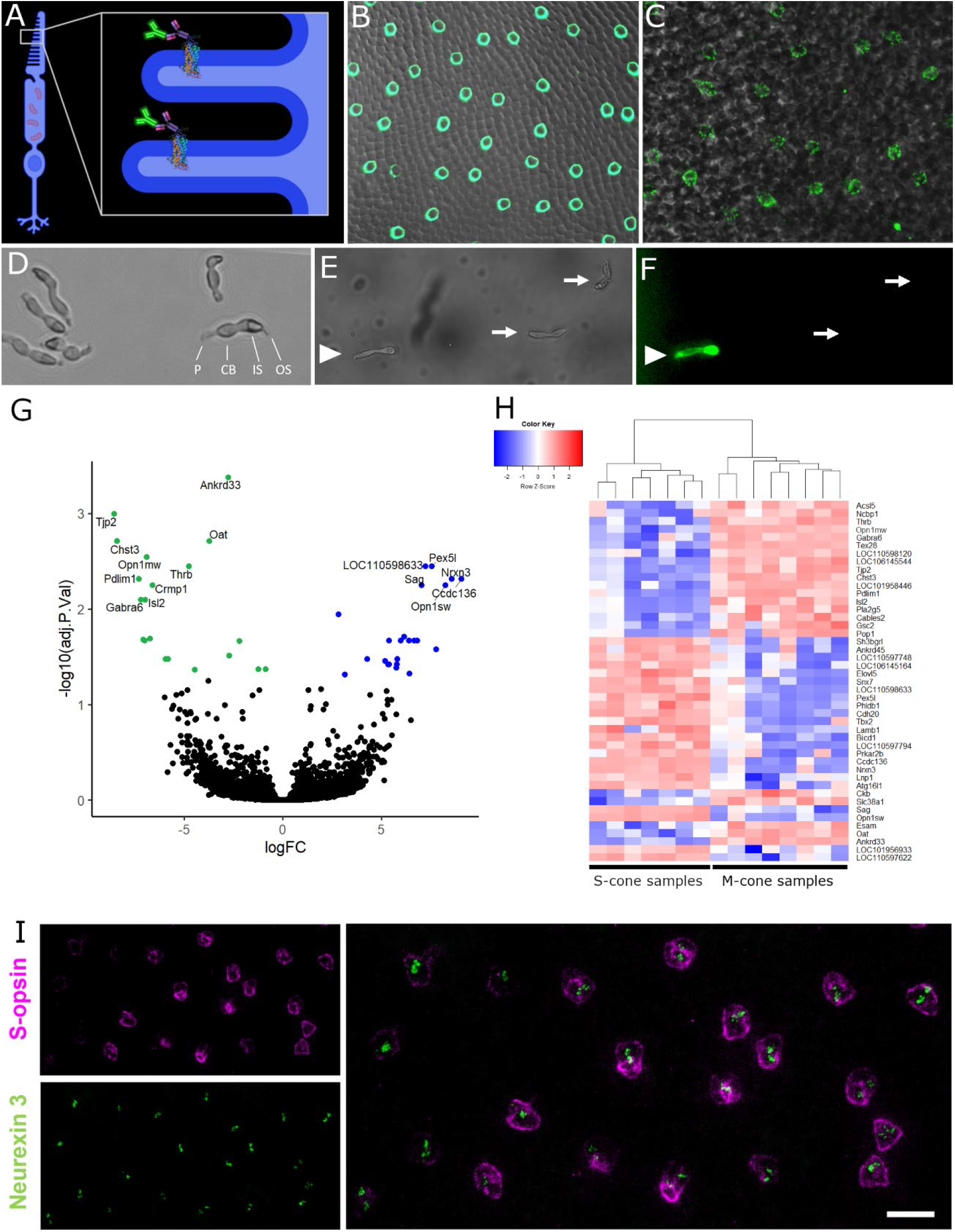
SMART-seq approach on manually collected M- and S-cone samples. A) An antibody targeting an extracellular epitope of S-opsin was used to label live S-cones. B) S-cone mosaic in a fixed ground squirrel retina preparation. C) Image of the S-cone mosaic in a live ground squirrel retina preparation. D) Cones after dissociation. P: pedicle, CB: cell body, IS: inner segment, OS: outer segment. E-F) Fluorescently labeled live cones after antibody incubation under (E) bright field and (F) epifluorescent lighting. Labeled cone (arrowhead) was categorized as S-cone, while cells with cone morphology but without label (arrows) were categorized as M-cones. G-H) Differentially expressed genes in M- and S-cone photoreceptors. G) Volcano plot highlighting genes significantly enriched in M-cones (logFC < 0) and S-cones (logFC > 0). H) Heat map showing expression of significant differentially expressed genes in all collected samples. I) Validation of cone subtype-specific gene expression of *Nrxn3*. Flat-mounted ground squirrel retina after single molecule fluorescent *in situ* hybridization (smFISH) with *Nrxn3*-probe, followed by S-opsin antibody-staining. Magenta: anti-S-opsin antibody. Green: *Nrxn3* smFISH probes. Scale bar: 10 μm.

Confirming the quality of our sequencing results, common cone genes such as *Gnat2, Cnga3, Cngb3* and *Grk7* were similarly expressed in both cone subtypes as expected, whereas the opsin genes *Opn1sw* and *Opn1mw* were selectively enriched in S-cones and M-cones, respectively. In addition, marker genes of other cell types, e.g., *Rbpms* (retinal ganglion cells), *Vim* (macroglia), *Calb1* (horizontal cells), and *Vsx2* (bipolar cells) were either not present or detected at low levels, confirming a lack of contamination by other cell types and thus indicating a high purity of the collected samples (Supplementary Table S1).

### The cell-adhesion molecule *Nrxn3* is specifically expressed in S-cones

We identified 58 genes that showed significant differential expression in S- or M-cones (adj. p-value < 0.1). Of these 58 DEGs, 28 genes were enriched in M-cones and 30 in S-cones (Figure 1 G-H). Filtering these DEGs for known cell adhesion moles, we identified S-cone specific expression of neurexin 3 (*Nrxn3*) as a potential candidate that may a play an important role at the S-cone to SCBC synapse.

Neurexins are cell-adhesion molecules involved in protein-protein interactions at synapses, where they form trans-synaptic complexes with many different protein families; e.g., neurexophilins, neuroligins, LRRTMs, latrophilins, and others (*22–29*). We first confirmed the specificity of *Nrxn3* via single molecule fluorescent *in situ* hybridization (smFISH) combined with antibody-based detection of S-opsin (Figure 1 I). The detected smFISH probes confirm that, within in the photoreceptor layer, *Nrxn3* expression is restricted to S-cones. Thus, we identified *Nrxn3* as a candidate molecule for involvement in the specific targeting of S-cones by SCBCs.

### *Nrxn3* is required for synaptic connections between S-cones and S-cone bipolar cells in mice

While sequencing of photoreceptors in the cone-dominant TLGS retina facilitated the identification of candidate genes for S-cone specific synaptic connections, the necessary genetic tools for confirming its function remain unavailable in TLGS. Thus, we proceeded with loss-of-function studies in the mouse, where such tools are readily available. To achieve S-cone specific ablation of *Nrxn3*, we crossed the Cre-dependent *Nrxn3* flox mouse, *Nrxn3*^tm3Sud^/J (*30*), with the S-opsin-Cre line Tg(Opn1sw-Cre). Cre recombinase is expressed under the control of the *Opn1sw*-promoter in about 12.3% of S-opsin-expressing cones. This provides cone-specific knockout of functional α- and β-neurexin 3 in those Cre^+^, S-opsin-expressing cones (S^+^/Cre^+^), leaving the remaining S-opsin-expressing cones (S^+^/Cre^-^) as wildtype controls. We then crossed these mice with a *Cpne9*-venus mouse line (*13*), which expresses the yellow fluorescent protein venus under the control of the *Cpne9*-promoter. *Cpne9* expression is highly specific for SCBCs, thus providing fluorescent labeling of SCBC dendrites and facilitating examination of contacts with S-cones (*31*). We then used opsin antibodies to identify “true” S-cones (those expressing S-opsin but lacking M-opsin; S^+^/M^-^), among which we differentiated those with normal *Nrxn3* expression (S^+^/M^-^/*Nrxn3*^*l*oxP/loxP^/Cre^-^) from those without *Nrxn3* (S^+^/M^-^ /*Nrxn3*^loxP/loxP^/Cre^+^). Additionally, we included Cre^+^ true S-cones from *Nrxn3*^+/+^ mice that retain *Nrxn3* (S^+^/M^-^/*Nrxn3*^+/+^/Cre^+^) as control true S-cones.

We then probed the phenotype of SCBC synaptic contacts for these different genotypes of true S-cones (Fig. 2). True S-cones with *Nrxn3* KO (S^+^/M^-^/*Nrxn3*^loxP/loxP^/Cre^+^) often lacked clear synaptic contacts with SCBCs, whereas such contacts were readily identified for control true S-cones (S^+^/M^-^/*Nrxn3*^*l*oxP/loxP^/Cre^-^ or S^+^/M^-^/*Nrxn3*^+/+^/Cre^+^, Fig. 2 B-C). To quantify the degree of normal true S-cone synapse disruption in an unbiased manner, we enlisted trained raters to score isolated anonymized confocal images of cone synaptic terminals from *Nrxn3* KO (*Nrxn3*^*l*oxP/loxP^/Cre^+^) and control (*Nrxn3*^+/+^/Cre^+^) mice (see Methods for details; (*32*)).

**Figure 2.**
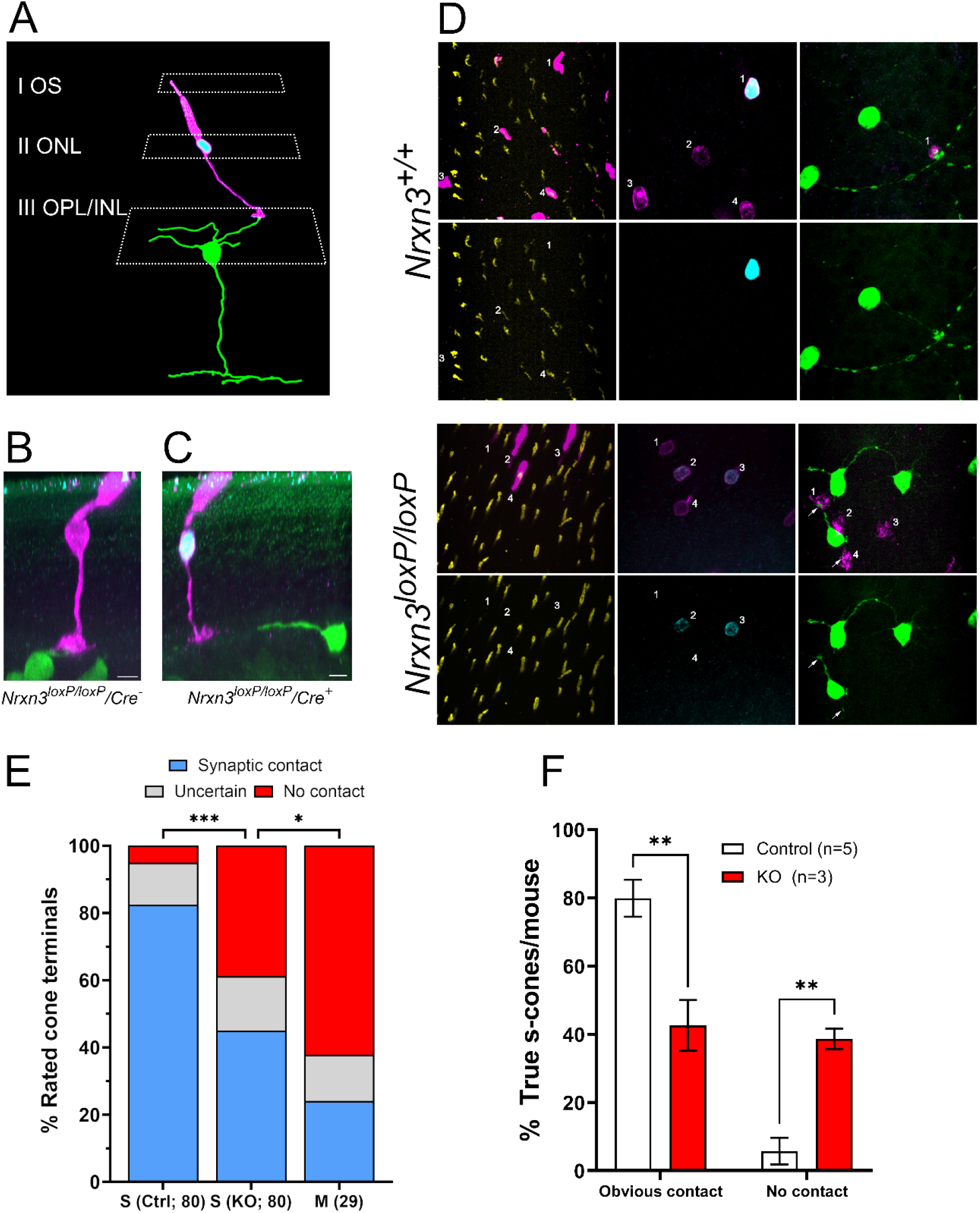
S-cone to S-cone bipolar cell (SCBC) synaptic connectivity in Nrxn3 KO mice. A) Method for classifying and quantifying cone-SCBC synapses. Cone identity was confirmed using S- and M-opsin antibody signals in the outer segment (OS, plane I). Next, the cell bodies of the outer nuclear layer (ONL) were imaged to determine Cre expression in cone cells (plane II). Lastly, cone pedicles and SCBC dendrites were imaged to identify contacts (plane III). All segments were acquired in a single confocal stack for later examination. B-C) Two representative S-cones from the same retinal sample, both of which are depth-corrected orthogonal projections from confocal Z-stacks acquired in the whole-mount configuration. B) a *Nrxn3* WT S-cone (*Nrxn3*^loxP/loxP^; Opn1sw-Cre^-^) with a normal SCBC contact. C) A *Nrxn3* KO S-cone (*Nrxn3*^loxP/loxP^; Opn1sw-Cre^+^) that fails to make contact with a nearby SCBC dendrite; Magenta: S-opsin, cyan: Cre, green: venus (SCBC). Scale bar: 5 μm. D) The upper panel shows S-cone to SCBC connectivity in a *Nrxn3* wild type control mouse. Cone 1 is a Cre^+^ S-cone which is contacted by two SCBCs in the same frame and potentially two further SCBCs which are out of frame. Cone 2 is a mixed cone (M-opsin^+^, S-opsin^+^) and Cre^-^. Nearby SCBCs do not contact this cone. The lower panel shows a *Nrxn3*^loxp/loxp^ sample in which Cre^+^ cells are lacking functional α- and β-neurexin 3. Cones 2 and 3 are both true S-cones (S^+^, M^-^) and express Cre. Nearby SCBCs appear to avoid these Nrxn3^-/-^ pedicles while pedicles from Cre^-^ control cones 1 and 4 appear to receive at least one SCBC-contact (arrows). Yellow: M-opsin, magenta: S-opsin, cyan: Cre, green: venus. E) Quantitative analysis of S-cone to SCBC contacts in control S-cones, *Nrxn3* KO S-cones, and mixed cones on a per-cone terminal basis; statistics computed as a two-proportions Z-Test comparing the proportions of terminals in each group rated as non-contacting. F) Rating proportions of contacts or no contacts on a per-animal basis, statistics computed using Student’s t-test. For all panels, statistical tests are indicated as follows: * = p<0.05; ** = p<0.01; *** = p<0.001.

Across the mice tested (3 *Nrxn3*^*l*oxP/loxP^/Cre^+^, 2 *Nrxn3*^+/+^/Cre^+^), a total of 189 cones were imaged and rated. Of the control true S-cones, 83% were rated as having obvious contacts with SCBCs (66 of 80), compared to 45% in *Nrxn3* KO true S-cones (36 of 80; Figure 2E). The increase in the proportion of SCBCs judged as failing to make contacts with true S-cones was statistically significant (p<0.001; two-proportions Z-test). Anatomically, mouse S-cone connectivity with SCBCs are not trivial to reliably identify even in wild-type retina (*33*). To account for false positives, e.g., *en passant* SCBC dendrites crossing near S-cone terminals, we also used mixed cones (S^+^/M^+^), which do not contact SCBCs (*13*), as a control for gauging this false positive rate. Such mixed cones were rated as contacting SCBCs in 24% of the images (7 of 29). On a per-animal basis (Fig. 2F), the proportion of cone terminals judged as clearly not forming a synapse with SCBCs was significantly higher (p<0.01; t-test) for true S-cones with *Nrxn3* KO (38.7 ± 3.0%, SEM; n=3) than that for control true S-cones (either S^+^/M^-^/*Nrxn3*^*l*oxP/loxP^/Cre^-^ or S^+^/M^-^/*Nrxn3*^+/+^/Cre^+^) (5.7 ± 3.9%, SEM; n=5). Thus, despite the uncertainty of the nature of the remaining contacts observed using confocal microscopy, we can conservatively estimate that more than 50% of true S-cone contacts with S-cone bipolar cells were eliminated in the *Nrxn3* KO mouse compared to *Nrxn3* WT control mice.

## Discussion

In this study, we effectively combined the strengths of both TLGS and mouse models to leverage their unique advantages. The TLGS model, characterized by its cone-dominance and the lack of mixed cones (those expressing multiple opsins), offered new insights into the gene expression specific to different cone subtypes. Meanwhile, the mouse models, empowered by their extensive genetic manipulation capabilities, allowed us to investigate the functional aspects. Through this synergistic approach, we were able to demonstrate the critical function of *Nrxn3* in facilitating and/or maintaining the precise connections between S-cones and their corresponding postsynaptic S-cone bipolar cells.

First, we acquired distinct genetic profiles between S- and M-cone photoreceptors and identified potential genes that may underlie critical differences in the wiring of these photoreceptors. Our findings suggest that *Nrxn3* is required for the first synapse in the S-cone pathway, evidenced by a significant reduction—down to 45%—in true S-cone contacts with SCBCs in *Nrxn3* knockout (KO) mice compared to wild-type counterparts. Prior research has primarily depicted *Nrxn3* as serving various regulatory functions at synapses. In studies of neurexins in the mammalian brain, the removal of one or more neurexin variants led to a range of outcomes, varying according to the brain region and specific combinations of pre- and postsynaptic partners. These outcomes include the assembly of presynaptic machinery (*22*), the stability of postsynaptic receptors (*30*), the plasticity of axonal processes, and a variety of synaptic activities in a neuronal subtype-specific manner (*23, 34*). Although neurexins are recognized for their extensive splice variants, our research is the first, to our knowledge, to identify a specific synapse within the central nervous system where *Nrxn3* is indispensable. It remains uncertain whether the reduction of S-cone to S-cone bipolar cell synapses in *Nrxn3* KO mice signifies a direct role of neurexin in cell-cell interaction or an indirect outcome of impaired synaptic functionality. Nevertheless, our discoveries contribute to understanding the varied roles of neurexins in the nervous system and demonstrate that *Nrxn3’s* specific expression in S-cones is crucial for S-cone synaptic connectivity in the outer plexiform of the retina.

### S-cone synaptic connections in the *Nrxn3* knock out mouse model

To investigate *Nrxn3’s* role in mammalian S-cones, we employed mouse models (*13, 30*). A recent study provided supporting evidence of a conserved link between S-cone identity and *Nrxn3* expression in mice: the deletion of Trβ2, a thyroid hormone receptor crucial for M-cone differentiation, led to reduced M-cone gene expression and increased S-opsin and *Nrxn3* expression in mouse cones, affirming the relevance of our mouse model (*35*). In our model, the Cre-dependent deletion of α- and β-neurexin 3 resulted in a loss of contacts with SCBC in approximately half of Cre^+^ true S-cones, indicating an important role for *Nrxn3* in this highly specific synapse. Based on previous studies (*8, 13, 15*), we would not expect SCBCs to form contacts with mixed cones (S- and M-opsin^+^). Nevertheless, our trained raters identified 24% of such cones as appearing to contact SCBCs. This finding could suggest non-specific baseline interactions between SCBCs and mixed cones; however, it is more likely to reflect inaccuracies by the raters, potentially identifying false positives. Notably, a closer look at mixed cones deemed ‘highly likely to contact SCBCs’ showed many resembled true S-cones with faint M-opsin signals, suggesting the classification of cone subtypes based on opsin expression in the mouse retina might be more graded than binary, as might be their connectivity patterns (Fig. S1). Regardless, deletion of *Nrxn3* significantly lowered the proportion of S-cones rated as making proper SCBC contacts, strongly supporting that *Nrxn3* deletion disrupts this specific wiring.

### Advantages and limitations of the *Nrxn3* KO model

In our study examining the effects of Nrxn3 loss in S-cones, we focused on young adult mice aged 60 to 70 days. As a result, it remains unclear whether the absence of neurexin 3 hinders the initial formation of synapses during development or affects their long-term maintenance. The failure to establish initial contacts could arise from S-cones not effectively attracting SCBC dendrites or from the inability to form successful synapses.

Future research will be necessary to explore the behavior of S-cone bipolar cell dendrites at earlier timepoints of retinal development to address these questions. Given the role of neural plasticity, it will also be crucial to determine whether there is an increase in contacts with other bipolar cell types and how such changes might influence color coding when S-cone to SCBC connections are diminished.

Additionally, the mosaic expression of Cre recombinase in our Opn1sw-Cre transgenic line allowed us to observe Cre^+^ Nrxn3-KO cone pedicles near Cre^-^ cones with intact *Nrxn3* expression within the same sample, as illustrated in Fig. 2 D. This setup was advantageous for directly comparing the morphological differences between normal and mutant S-cone contacts. However, to assess the impact of disrupting the first synapse in the S-cone pathway on mouse behavior and overall light responses, a different Cre line that ensures a complete loss of *Nrxn3* in all S-cones will be necessary.

### Deep sequencing of isolated thirteen-lined ground squirrel photoreceptors

In our study, we employed the SMART-seq technique to explore the differential gene expression among cone subtypes, contrasting with high-throughput droplet-based sequencing methods (*20*). The success of this approach largely hinged upon the usage of TLGS, for three main reasons: first, they have a cone-dominant retina. Second, their S- and M-cones are clearly distinguishable since there is no co-expression of M- and S-opsin in the same cell. Third, squirrel neurons are more resilient to damage and provide higher quality material as they showed constantly high cell viability of > 92% after dissociation (determined with propidium iodide staining; data not shown). We speculate this could be due to the cell-protective mechanisms present in this hibernating animal, which has the genetic programing to withstand extreme conditions (*36, 37*). The combination of these features enabled us to generate high-quality, low-contamination genetic profiles for S- and M-cones, and consequently a more complete list of differentially DEGs compared to studies that utilized high-throughput droplet-based sequencing protocols (*18, 19*). While the droplet-based sequencing method is ideal for classifying large amounts of cell types, this single-cell collection-based SMART-seq method enhanced the detection of genes with lower expression levels such as *Nrxn3*, which has not been previously identified as a DEG in S-cones.

Thus, our dataset provides a valuable roadmap for exploring the functions of these DEGs and their cone subtype-specific physiology. For example, we identified higher expression levels of rod arrestin (*Sag*) in S-cones, but not in M-cones, while cone arrestin (*Arr3*) exhibited the opposite pattern (Fig.1 G-H). This may underly the difference in the kinetics of S- and M-cone light response, especially the decay phase to which the arrestin molecules contribute (*38*). Conversely, the gene *Tjp2*, specific to M-cones, encodes the tight junction protein 2 that may be responsible for the M-cone-specific cell membrane localization of connexin 36 (*39*), which ensures electrical coupling exclusively between M-cones. Taken together, our results highlight the advantage of SMART-seq when investigating a small number of specialized cells, such as S-cones. This strategic approach will be invaluable for targeting postsynaptic SCBCs and to identify the postsynaptic binding partner(s) of neurexin 3. Such insights are fundamental in unraveling the synaptic formation processes in the S-cone pathway. Understanding these molecular mechanisms for proper synaptic connections is essential for the advancement of cell replacement therapies for the retina, offering potential therapeutic avenues for individuals with visual impairments.

## Materials and Methods

### SMART-seq of ground squirrel photoreceptors

Eyes were enucleated from euthanized ground squirrels by sharp dissection following preparation of the retina and removal of the retinal pigment epithelium. A ∼10 mm^2^ piece of retina was incubated in digestion buffer (5 U/mL papain, 0.667 mg/mL L-cysteine, 1 mM EDTA in Hank’s balanced salt solution (HBSS)) for 20 min at 37 °C. The sample was washed two times with enriched Hibernate-A media (Hibernate-A medium, 2% B27, 1 % streptavidin/penicillin, 0.25 % Glutamax, 0.25 % L-glutamine, 10 mM NaCl) followed by antibody incubation (anti-S-opsin) for 45 min (4 °C). After two washes, the retina was incubated with Donkey-anti-goat antibody for 45 min at 4 °C. The tissue was triturated 10 times using a 1000 µL pipette tip to dissociate cells. The single cell suspension was filtered through a cell strainer (35 µm) and centrifuged to remove debris (2000 x g, 3 min, 4 °C). The pellet was carefully resuspended in 500 µL enriched Hibernate-A and placed on an inverted microscope (Evos cell imaging system). S-opsin-positive cells (S-cones) and negative cells with cone morphology (M-cones) were separately collected using a micromanipulator (Eppendorf TransferMan 4r) and transferred into 8 µL lysis buffer (Takara Bio SMARTer Ultra Low Input RNA Kit for Sequencing - v3). For each sample, 16–20 single cells were pooled. Samples were flash frozen in dry ice and temporarily stored at -80 °C until RNA extraction and library preparation. RNA-extraction of single cell samples and generation of double stranded cDNA was performed using the Takara Bio SMARTer Ultra Low Input RNA Kit. For the construction of the library, the Low Input Library Prep Kit v2 (Takara Bio) was used. cDNA quality was analyzed with the Agilent 2100 BioAnalyzer and concentrations were measured with the Qubit dsDNA HS assay. Up to eight samples with different Illumina barcodes were pooled in one lane of a flow cell (Illumina HiSeq 2500; 50 bp read length, single-end mode). All kits were used as instructed by the manufacturers.

### RNA-seq data analysis

Analysis was performed as described (*40*). Briefly, we used RefSeq transcriptome and annotation GCF_000236235 to estimate abundance of ground squirrel transcripts using the Kallisto package (*41*) on the NIH high performance computer Biowulf cluster. Resulting output was imported into R (*42*) using the packages rhdf5 (*43*) and tximport (*44*) and then normalized and analyzed for differential gene expression using R packages EdgeR (*45*) and limma (*46*).

### Immunohistochemistry and image acquisition

Dissected retinae were incubated in PBS with 4 % PFA (mouse: 30 min, squirrel: 45 min) at room temperature (RT). After three washes with PBS for 15 min, the tissue was incubated with primary antibody (Table 1) in blocking solution (PBS with 4 % normal donkey serum, 0.01 % sodium azide, 0.5 % Triton X-100) for five days at 4 °C. After washing five times (15 min) with PBST (PBS + 0.5 % Triton X-100), the tissue was incubated with secondary antibody (1:500) in blocking solution for two days at 4 °C. The samples were washed 5 times with PBST before mounting on a microscope slide. Retinal whole-mounts were imaged with a 60x objective on a Nikon A1R confocal microscope controlled by Nikon NIS-Elements software or a 63x objective on a Zeiss LSM 780 microscope controlled by Zeiss Zen software.

**Table 1.** Antibodies used in this study. IF: immunofluorescence.

| Antibody | Source | Catalog number | Dilution |
| --- | --- | --- | --- |
| Anti-GFP (chicken) | Aves labs | GFP-1020 | IF: 1:100 |
| Anti-S-opsin (goat) | Santa Cruz | sc-14363 | IF: 1:200;<br>Picking: 1:50 |
| Anti-Cre (guinea pig) | Synaptic Systems | 257 004 | IF: 1:250 |
| Anti-M/L-opsin (rabbit) | Millipore | AB5405 | IF: 1:200 |
| Anti-chicken Alexa 488 | Jackson ImmunoResearch | 703-545-155 | IF: 1:500 |
| Anti-goat Alexa 647 | Jackson ImmunoResearch | 705-606-147 | IF: 1:500 |
| Anti-goat Alexa 488 | ThermoFisher | A-11055 | Picking: 1:50 |
| Anti-guinea pig Cy3 | Jackson ImmunoResearch | 706-165-148 | IF: 1:500 |
| Anti-rabbit DyLight 405 | Jackson ImmunoResearch | 711-475-152 | IF: 1:500 |

### Oligopaints smFISH (single molecule fluorescent *in situ* hybridization) probe design and synthesis

SmFISH probes were designed for Ictidomys tridecemlineatus neurexin 3 (*Nrxn3*) transcript variant X1 (XM_021723849, NCBI database) using the OligoArray software package (*47*) with a target length of 30 bp, which resulted in 140 unique probe sequences covering the full length of the *Nrxn3* transcript. Custom primer sequences were appended to the 5’ and 3’ ends of all *Nrxn3* probes prior to synthesis as part of a complex ssDNA library (CustomArray Inc.). *Nrxn3* probes were PCR amplified from the ssDNA library, concentrated via column purification, and labeled with fluorophores in a second PCR reaction using fluorophore-labeled PCR primers (IDT). Fluorescent dsDNA probes were then converted to ssDNA via Lambda Exonuclease digestion, and concentrated by ethanol precipitation as described (*48*).

### SmFISH procedure

Squirrel retinae were dissected in nuclease-free PBS and cut into rectangular pieces (approx. 2 mm x 3 mm), followed by fixation in 4 % PFA in PBS for 15 min at RT. After three washes with PBS, the tissue was permeabilized with 0.5 % Triton X-100 in PBS for 30 min (rocking). Tissue was washed with 2X saline-sodium citrate (SSC) buffer containing 10 % formamide for 10 min (rocking). Tissue was incubated with probe mix (10 μl of *Nrxn3* smFISH probe + 90 μl hybridization buffer (30 % formamide, 10 % dextran sulfate, 2 mM vanadyl ribonucleoside complex in SSC buffer)) over night at 37 °C on a shaking incubator. The sample was briefly rinsed and then washed for 30 min with 2XSSC and 30 % formamide at 37 °C. The sample was then briefly rinsed, and two times washed for 30 min with 2XSSC and 10 % formamide at 37 °C. After a final rinse with 2XSSC, the tissue was either mounted and imaged or subjected to subsequent immunohistochemical staining for co-labeling with S-opsin antibody.

### Mouse strains

The transgenic *Opn1sw*-Cre mouse was generated by the NEI Genetic Engineering Core by BAC insertion. Strain generation was identical to that of our previously described *Opn1sw*-venus mouse (*49*), but with the Cre recombinase sequence instead of venus. Cre expression is under control of the *Opn1sw*-promoter. To quantify the proportion of Cre-expressing S-opsin^+^ cones, we imaged retinal flat-mounts from three *Opn1sw*-Cre mice after staining with S-opsin- and Cre-antibody and corresponding fluorescent secondary antibodies. On average, 12.3% of S-opsin^+^ cells also expressed Cre. The S-cone bipolar cell *Cpne9*-venus mouse was previously described in Nadal-Nicolás et al. (2020). The *Nrxn3*^tm3Sud^/J mouse was generated by the Südhof lab (*30*) and obtained from The Jackson Laboratory (strain #014157), and backcrossed over multiple generations to C57BL/6J mice (strain #000664). Mice in this study were both female and male, 60-70 days old. All experiments and animal care were conducted in accordance with protocols approved by the Animal Care and Use Committee of the National Institutes of Health.

### Imaging rating and statistical analysis

From each mouse (5 total: 3 *Nrxn3* KO: *Nrxn3*^loxP/loxP^; Tg(Opn1sw-Cre^+^) and 2 *Nrxn3* wild type: *Nrxn3*^+/+;^ Tg(Opn1sw-Cre^+^)), confocal image stacks of S-opsin^+^ cones were acquired, and each cone was categorized as a “true” S-cone vs. a mixed cone (using the presence vs. absence of M-opsin labeling in the cone outer segment). S-cones were further divided into *Nrxn3* KO vs. control (according to the Cre antibody signal in the cone nucleus). From the acquired z-stack images, sections of the image surrounding the synaptic terminal were extracted that contained only the S-opsin and venus labels (marking SCBCs), thus isolating context-free images of the S-cone terminal and S-cone bipolar cell dendrites. Cone categorization was confirmed independently by two researchers.

These images were assigned randomized numbers to anonymize the cone categorization and were then shared with volunteer raters (2 participants; see also Hallgren et al. (*32*)) who were first trained on the task—without having been informed of the expected outcome of this analysis—and then asked to score each terminal from 1 (“clearly no contact”) to 5 (“a clear synaptic connection”). Raters were experienced with using ImageJ to view 3D multichannel confocal z-stacks for judging each terminal. Completed ratings were returned to a third expert rater, who noted the rankings for each anonymized image. Final ratings for each image were taken as the average of the two raters’ scores, or in the case of major disagreements (defined as score differences of 2 points or more), the expert rater assigned a replacement score to the still-anonymized images based on their own judgment (21 of 195 images). For each mouse, the proportions of S-cone images scored higher than 3.5 (scores of 4 or 5 were described as “obvious contact” and “contact likely” respectively) were computed for KO vs. control S-cones and compared (Fig. 2F). KO S-cones were defined as M-opsin^−^, Cre^+^ cones in Nrxn3^loxP/loxP^ mice, and control S-cones were defined as either M-opsin^−^, Cre^−^ cones in Nrxn3^loxP/loxP^ mice, or M-opsin^−^ cones in wild-type mice (*Nrxn3*^+/+^) regardless of Cre labeling. Observed proportions of these highly-rated S-cone images were compared using a one-tailed t-test using Microsoft Excel. Statistical tests on proportions of all images (Fig. 2E) were computed upon the proportions of images across conditions rated lower than 2.5 (scores of 1 and 2 were described as “clearly no contact” and “contact unlikely” respectively) using a two proportions Z-test. For all statistical tests, significance was denoted as follows: *: p<0.05, **: p<0.01, ***: p<0.001.

## Acknowledgments

We thank Christian Puller, Gui-shuang Ying and Douglas Forrest for helpful ideas and discussions. We would like to thank the NIH animal facility staff for year-round care of our animals.

## Author contributions

Conceptualization: VPK, JMA, WL

Methodology: VPK, JA, JMB, MBT, FMN

Investigation: VPK

Image rating: XL, AS

Formal analysis: VPK, JMB

Visualization: VPK, JMB

Supervision: WL

Writing—original draft: VPK

Writing—review & editing: VPK, JMB, WL

## Supplementary Materials

**Table S1.** Expression values (counts per million) of marker genes in S- and M-cone samples. Low number of marker genes from other cell types indicate little to no contamination. Cone genes were detected as expected.

| Gene | Average expression in counts per million (cpm) after TMM<br>(trimmed mean of M values) normalization |  |  |  | Expected cell type |
| --- | --- | --- | --- | --- | --- |
|  | Combined S-cone samples |  | Combined M-cone samples |  |  |
| *Opn1sw* | 20681.26 | ± 6525.83 | 433.94 | ± 856.57 | S-cones |
| *Opn1mw* | 11.48 | ± 11.75 | 740.23 | ± 186.90 | M-Cones |
| *Cnga3* | 95.22 | ± 54.65 | 92.33 | ± 73.28 | All cones |
| *Cngb3* | 1618.16 | ± 457.70 | 1249.61 | ± 251.12 | All cones |
| *Gnat2* | 2837.57 | ± 929.58 | 3905.72 | ± 777.66 | All cones |
| *Grk7* | 400.83 | ± 195.09 | 460.50 | ± 126.99 | All cones |
| *Cnga1* | 5.28 | ± 12.86 | 14.23 | ± 37.66 | Rods |
| *Cngb1* | 112.44 | ± 61.80 | 25.05 | ± 20.40 | Rods |
| *Gnat1* | 0.00 | ± 0.00 | 0.02 | ± 0.06 | Rods |
| *Grk1* | 3.11 | ± 5.58 | 2.46 | ± 3.85 | Rods |
| *Rho* | 34.65 | ± 66.66 | 33.61 | ± 78.51 | Rods |
| *Gfap* | 0 | ± 0 | 0.00 | ± 0 | Astrocytes |
| *Vsx2* | 3.18 | ± 7.80 | 0.06 | ± 0.11 | Bipolar cells |
| *Rbpms* | 0.061 | ± 0.15 | 0.90 | ± 2.30 | Ganglion cells |
| *Pou4f2* | 0 | ± 0 | 0.00 | ± 0 | Ganglion cells |
| *Calb1* | 86.13 | ± 137.52 | 38.38 | ± 34.67 | Horizontal cells |
| *Vim* | 17.49 | ± 21.94 | 12.95 | ± 12.76 | Müller Glia |

**Fig. S1.**
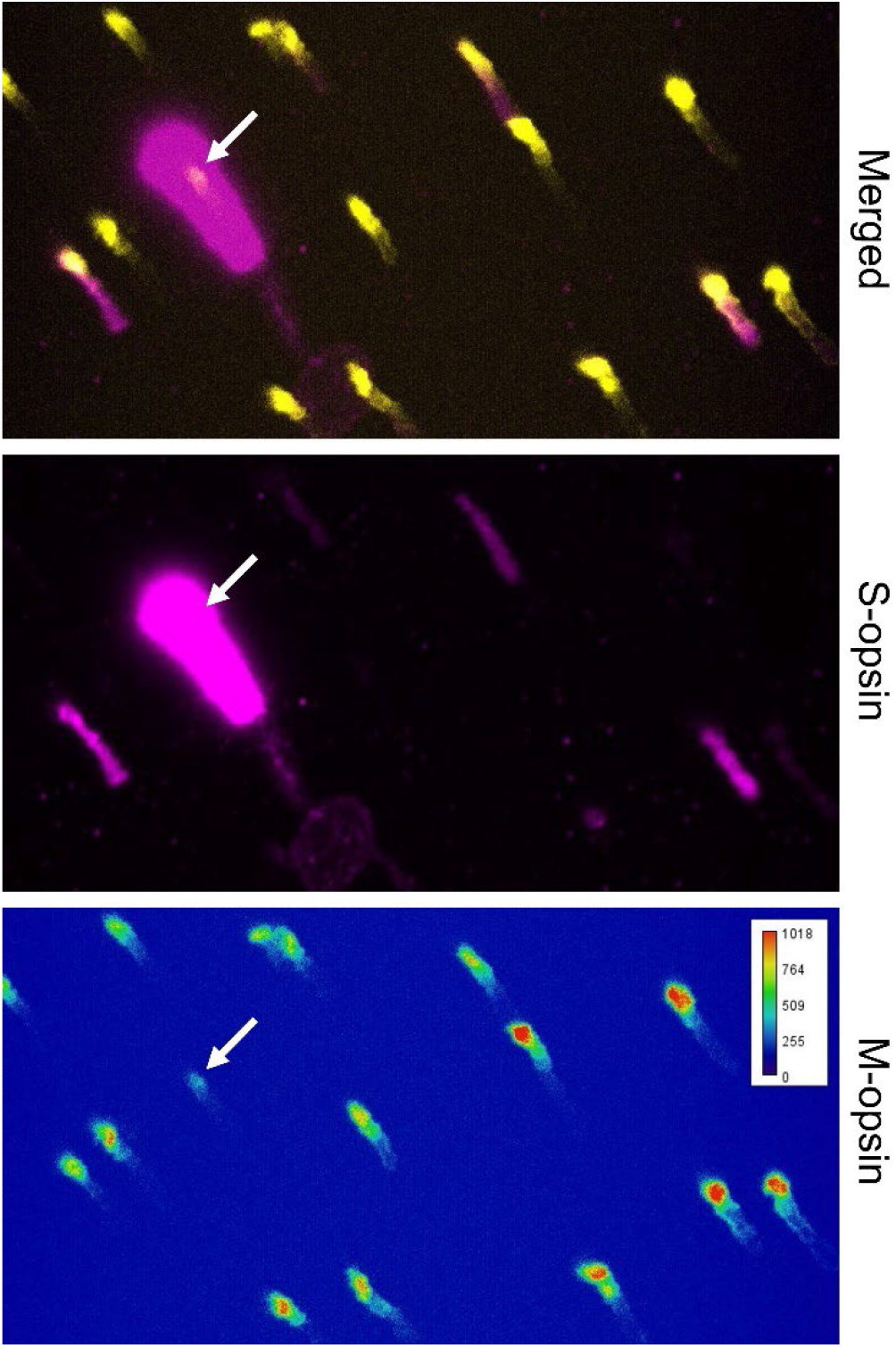
This image shows a cone (arrow) that received a clear SCBC contact (not shown here) but was categorized as mixed cone, because its outer segment is S-opsin^+^ and M-opsin^+^. For this cone, the strength of the M-opsin signal appears lower, and the S-opsin signal appears higher compared to surrounding M-opsin^+^/S-opsin^-^ or M-opsin^+^/ S-opsin^+^ cones, none of which were contacted by SCBCs.

